# SanA is an inner membrane protein mediating the early stages of Salmonella infection

**DOI:** 10.1101/2024.01.05.574334

**Authors:** Adrianna Aleksandrowicz, Rafał Kolenda, Teresa L M Thurston, Krzysztof Grzymajło

## Abstract

Bacterial membrane proteins, crucial for the interaction with the environment, encompass various functional molecules such as SanA. SanA is pivotal for the physicochemical properties of the bacterial membrane, influencing *Salmonella*’s antibiotic resistance and infection phenotype. Previous studies identified a link between *sanA* mutation and increased *Salmonella* invasiveness, but the mechanisms underlying this phenomenon remain largely unexplored. Therefore, our research investigates SanA’s role during *Salmonella* infection, examining its expression pattern, localization within the cell, and association with *Salmonella* Pathogenicity Island I (SPI-I). Using subcellular fractionation and Western Blotting we revealed that SanA is predominantly located in the inner membrane. Additionally, we utilized transcriptional fusion to monitor SanA expression under various environmental conditions. We observed that SanA plays a significant role during the late exponential and early stationary growth phase and remains important 24 hours after the bacteria enter host cells. Moreover, our invasion assays demonstrated that deletion of *sanA* in bacteria grown to early stationary phase significantly enhances their invasiveness, partly due to increased SPI-I expression, which is regulated in a nutrient availability-dependent manner. Our results highlight SanA’s essential role in *Salmonella*’s response to environmental stress, critical for its entry and survival in hostile environments. This research underscores the importance of inner membrane proteins in bacterial pathogenicity, particularly in the initial stages of infection.

## Introduction

*Salmonella enterica* stands out as one of the most prominent bacterial pathogens, causing food-borne diseases with significant morbidity and mortality in both humans and livestock (Ferrari et al., 2019). A critical stage in *Salmonella*’s pathogenicity depends on its ability to adhere to and invade host cells (Pizarro-Cerdá and Cossart, 2006). The bacterial structures employed in these processes vary widely, extending from monomeric proteins and multimeric macromolecules to intricate molecular machines. Notable for *Salmonella*, specialized type III secretion system (T3SS) encoded by genes of *Salmonella* Pathogenicity island I (SPI-I) and II (SPI-II) facilitate bacteria to invade and survive within phagocytic and non-phagocytic cells. It includes mechanisms that involve the translocation of the effector proteins into the host cell, thereby altering vesicular trafficking and cytoskeletal dynamics (Coburn et al., 2007).

*Salmonella* evades the host’s intracellular immune responses and persists in adverse environments by developing a sophisticated and complex cell envelope, which not only offers protection but also facilitates the selective passage of nutrients from the outside and removal of waste products from the inside (Silhavy et al., 2010). The envelope bilayer, composed of an outer membrane (OM) and an inner membrane (IM), showcases complexity and versatility, highlighted by numerous embedded proteins, each playing a specialized role. The OM with tightly packed lipopolysaccharide (LPS), provides a range of outer membrane proteins (OMPs) and functions as both a selective barrier and a platform for contact with the external environment (Silhavy et al., 2010). OMPs, including porins and efflux pumps, balance nutrient uptake and toxin exclusion, thus ensuring cellular homeostasis and protecting against threats like xenobiotics by moderating their intracellular accumulation (Sun et al., 2022). On the other hand, the IM, although less directly exposed to extracellular environments, harbors inner membrane proteins (IMPs) that participate in vital cellular processes, such as ATP synthesis and nutrient translocation (Silhavy et al., 2010). The synergistic interplay between OMPs and IMPs complexes constitutes a key adaptive mechanism in bacterial response to environmental stressors. These interactions strengthen the bacterial cell’s permeability barrier, enhancing resistance to xenobiotics and antimicrobial agents (Boughner and Doerrler, 2012).

Given the diverse functionalities of the bacterial envelope, Gram-negative pathogens utilize various modifications to the membranes, aiming to enhance their resilience to environmental stress and to successfully establish infections. For instance, it was shown that *Enterobacteriaceae* decrease porin expression as a quick response to toxic agents (Dam et al., 2018). Additionally, they modify lipopolysaccharides (LPS) to change the characteristics of the outer membrane, which helps in avoiding recognition by the host’s immune system and enhances their resilience to antimicrobial peptides. These modifications include altering lipid A phosphates, the core oligosaccharide phosphates, and lipid A acylation (Simpson and Trent, 2019). Adding to the complexity, other membrane attributes, like charge and hydrophobicity, modulate bacterial resistance to external stresses and were shown to affect pathogenicity indirectly. It was demonstrated that the efficiency of phagocytosis increases with the hydrophobicity of bacterial cells and that hydrophilic bacteria resist ingestion by phagocytes (Matz and Jürgens, 2001).

The link between membrane permeability and pathogenicity is further highlighted in *Salmonella*. Many of *Salmonella*’s invasion factors, such as T3SS-1, flagella, and chemotactic receptors, are integral components of its envelope. This association suggests a complex interplay between membrane permeability and the expression of virulence factors. Previous studies demonstrated that the expression of *hilD*, a principal regulator of the SPI-I, not only increases membrane permeability but also makes *Salmonella* more susceptible to membrane stress (Sobota et al., 2022). It highlights that the expression of virulence genes, while critical for pathogenesis, can also impose a significant fitness cost on pathogenic bacteria to maintain the balance between virulence and survival. Our current research has spotlighted the role of SanA in modulating the properties of the bacterial membrane, influencing its charge and hydrophobicity, which in turn affects antibiotic resistance and enhances the intracellular survival of *S.* Typhimurium (Aleksandrowicz et al., 2024). Additionally, our previous work demonstrated that a 10-nucleotide deletion in the *sanA* coding sequence is associated with enhanced invasive capabilities of bacteria (Kolenda et al., 2021). However, the mechanism underlying this phenotype has not yet been identified.

In light of these findings, this study aims to conduct a thorough examination of SanA’s expression during infection, its subcellular localization, and its interplay with SPI-1. Furthermore, we delve into SanA’s influence on the infection process of *Salmonella in vitro*.

## Material and Methods

### 1. Bacteria, plasmids, and growth conditions

**Tables 1**, **2**, and **3** contain the complete list of bacterial strains, plasmids, and primers used in this study, respectively. All the *Salmonella* strains employed were derived from the *Salmonella* enterica serovar Typhimurium 4/74 wild type (WT). Unless specified otherwise, bacterial cultures were typically cultivated at 37°C for 16 h (overnight) either in Lysogeny Broth (LB) under dynamic conditions with shaking (180 rpm), or on agar plates. For all invasion studies, *Salmonella* strains were grown under SPI-I-inducing conditions (early stationary growth phase in LB medium; OD_600_=2.0). When activation of SPI-II was required, the bacterial strains were grown overnight in LB medium and then washed in Mg-MES minimal medium (consisted of: 170 mM 2-(Nmorpholino)ethanesulfonic acid (MES) at pH 5.0, 5 mM KCl, 7.5 mM (NH_4_)_2_SO_4_, 0.5 mM K_2_SO_4_, 1 mM KH_2_PO_4_, 10 mM MgCl_2_, 38 mM glycerol, and 0.1 % Casamino Acids), with the pH adjusted to 5.0. The bacteria were then grown for 6 h in the same medium.

If required, antibiotics were supplemented at specific concentrations: 100 μg/ml for Ampicillin (Amp); 50 μg/ml for Kanamycin (Km), and 500 µg/ml for Erythromycin (Ery). For inducing *lac* and *ara* promoters, isopropylthio-β-galactoside (IPTG) was added to a final concentration of 0.5 mM or arabinose to a final concentration of 0.2 %, respectively. Cell growth was analyzed using optical density readings at 600 nm.

### 2. Cell culture

Human intestinal epithelial cell line Caco-2 (DMSZ, Germany) was grown at 37°C with 5 % CO_2_ in Dulbecco’s Modified Eagle’s Medium (DMEM)/Ham’s F-12 supplemented with 1 mM l-glutamine, 100 U/ml penicillin-streptomycin and 10 % fetal bovine serum (FBS) and passaged in the log phase of growth (at a confluency of 80-90 %) according to standard protocols. For infection assays cells were seeded in 24-well plates at a density of 1.2x10^5^ cells and used in experiments after five, six, or seven days.

Immortalized bone marrow-derived macrophages (iBMDM) were maintained in DMEM high glucose supplemented with 20 % (vol/vol) of L929-MCSF supernatant (LCM), 10 % (vol/vol) of FBS 10 mM of HEPES, 1 mM of sodium pyruvate, 0.05 mM of β-mercaptoethanol and 100 U/ml of penicillin/streptomycin and seeded at a concentration of 1x10^6^ or 2x10^5^ cells per well in a 6-well or 24-well plate, respectively 24 h prior infection.

### 3. Growth curves

To determine the growth curves, LB medium was inoculated with an individual bacterial colony and incubated overnight at 37°C with agitation at 180 rpm. The resulting cultures were then diluted to an optical density (OD_600_) of 0.05 in LB medium and incubated until the early log-phase growth (OD_600_=0.5) at 37°C, 220 rpm. Subsequently, cultures were centrifuged, rinsed with 0.9 % NaCl, and resuspended in the same solution. Optical density was assessed and cultures were further diluted in LB medium to achieve a bacterial concentration of 5x10^6^ CFU/ml. Optical density measurements were taken in Tecan Spark Control (Tecan) at 15-minute intervals over a 16 h period, and the cultures were shaken 30 seconds before each measurement. The study was conducted with a minimum of three independent biological replicates, and dilution series were set up on LB agar plates for verification of the initial bacteria number.

### 4. Cloning of *sicA* and its promoter

All genes or their promoters were amplified from *S*. Typhimurium 4/74 strain. Amplification was carried out by PCR using Phusion polymerase (Thermo) with primers listed in **Table 3**. PCR products were purified by GeneJET PCR purification kit (Thermo) whereas plasmid DNA was isolated by GeneJET Plasmid Miniprep Kit (Thermo). For creating a dual reporter with constitutive mCherry expression and inducible GFP expression, *sicA* promoter was inserted into pFCcGi plasmid in MluI/XbaI digestion sites. For HA-based reporter, *sicA* with promoter sequences were cloned into pFPV25.1GFPmut3.1Kan-2xHA in SacI/BglII digestion sites. DNA sequence of all the inserts was confirmed by Sanger sequencing.

**Table 1.** Bacterial strains used in this study.

| Strain | Relevant feature(s) | Reference |
| --- | --- | --- |
| *Salmonella enterica* subsp. *enterica* serovar Typhimurium 4/74 | wild type (WT) | (Aleksandrowicz et al., 2024) |
| *S. Typhimurium* *sanA*<sub>RBS::luc</sub> | *sanA*<sub>RBS::luc</sub> | This study |
| *S. Typhimurium* pFCcGi-*psicA* | pFCcGi-*psicA* | This study |
| *S. Typhimurium* pFPV25.1GFPmut3Kan-*sicA*-2xHA | pFPV25.1GFPmut3Kan-*sicA*-2xHA | This study |
| *S. Typhimurium* 4/74 Δ*sanA* | *S. Typhimurium* 4/74 with *sanA* gene knockout | (Aleksandrowicz et al., 2024) |
| *S. Typhimurium* Δ*sanA* pWSK29 | pWSK29 | (Aleksandrowicz et al., 2024) |
| *S. Typhimurium* Δ*sanA* pWSK29-*sanA* | pWSK29-*sanA* | (Aleksandrowicz et al., 2024) |
| *S. Typhimurium* Δ*sanA* pFCcGi-*psicA* | pFCcGi-*psicA* | This study |
| *S. Typhimurium* Δ*sanA* pFPV25.1GFPmut3Kan-*sicA*-2xHA | pFPV25.1GFPmut3Kan-*sicA*-2xHA | This study |
| *E. coli* XL1-Blue | *recA1 endA1 gyrA96 thi-1 hsdR17 supE44 relA1 lac [F'* | Wroclaw University of Environmental and |
|  | *proAB lacIq Z<sup>-</sup>M15 Tn10 (Tetr)* | Life Sciences,<br>Department of<br>Biochemistry and<br>Molecular Biology<br>collection |

**Table 2.** Plasmids used in this study.

| Plasmid | Relevant feature(s) | Reference |
| --- | --- | --- |
| p3121 | *luc* template vector, AmpR | (Gerlach et al., 2007) |
| pFCcGi | vector with arabinose-inducible expression of GFP and constitutive expression of mCherry, AmpR | (Figueira et al., 2013) |
| pFC-*psicA* | based on pFCcGi; with constitutive mCherry expression and GFP expression under the control of *sicA* promoter, AmpR | This study |
| pFPV25.1GFPmut3Kan | plasmid with GFPmut3 under the constitutive *rpsM* promoter, multiple cloning site and hemagglutinin tag (HA), KanR | Dr Rafał Kolenda,<br>Department of<br>Biochemistry and<br>Molecular Biology,<br>Wrocław University of<br>Environmental and Life<br>Sciences, Wrocław,<br>Poland |
| pFPV25.1GFPmut3Kan-*sicA*-2xHA | pFPV25.1GFPmut3Kan vector with *sicA* and its promoter insert, KanR | This study |

**Table 3.**
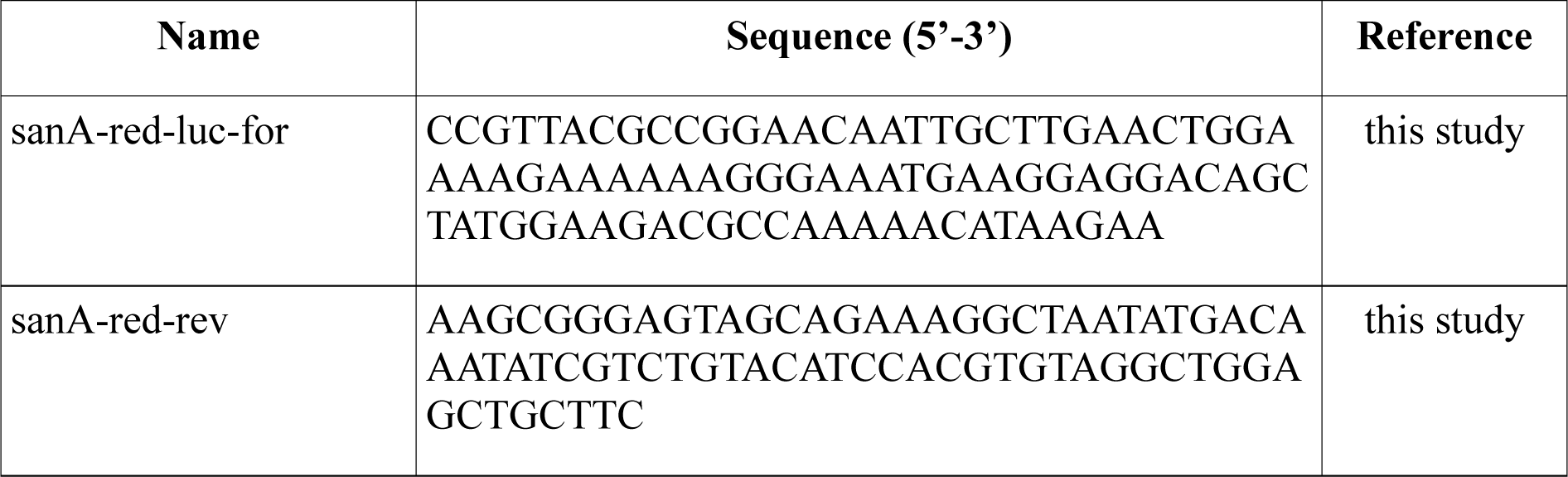
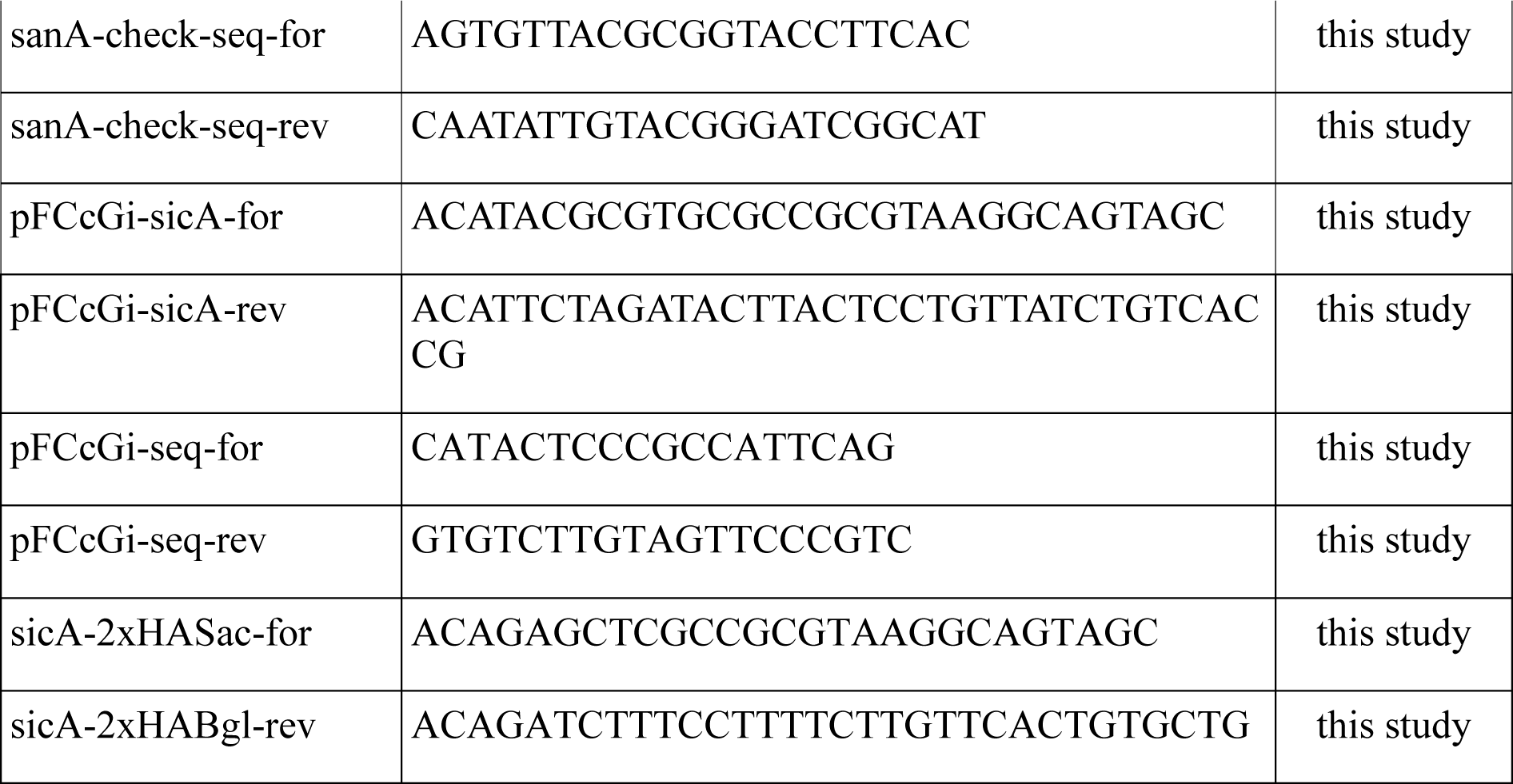
Primers used in this study.

### 5. Infection assay

Bacteria were grown under conditions optimizing SPI-I-dependent invasion or SPI-II-dependent replication within macrophages. For SPI-I-inducing conditions, an overnight culture was subcultured in LB at 37°C until the early stationary growth phase (OD_600_=2.0) (Peterson J. W, 1996). For SPI-II-inducing conditions bacteria were grown until the late stationary growth phase (overnight culture) (Martínez et al., 2014). iBMDM monolayers were infected with stationary phase bacteria opsonized in mouse serum for 20 min using a multiplicity of infection (MOI) of 10:1. To synchronize the infection, the plates were centrifuged for 5 min at 165 x g, followed by a 30-min incubation at 37°C (5 % CO_2_). Fresh DMEM supplemented with 100 µg/ml of gentamicin (Gm) was added to kill extracellular bacteria, and the macrophage monolayers were incubated with added Gm for 90 minutes (Monack et al., 1996). After washing with DMEM, the monolayers were lysed in 1 % Triton X-100 and diluted with PBS. Dilutions of the suspension were then plated on LB agar medium to assess the number of viable bacteria. To evaluate intracellular growth, the medium containing 100 µg/ml Gm was replaced with DMEM supplemented with 10 µg/ml of Gm, and parallel cell cultures were examined for viable bacteria 24 h following infection. Similarly, for the Caco-2 invasion assay, early stationary phase bacteria were added to the monolayer until the final MOI=100, cells were treated as described above and CFU/ml of bacteria was determined by plating dilutions of the suspension on LB agar.

### 6. Determination of SanA expression during infection

Transcriptional fusion was created as described previously by Gerlach et al (Gerlach et al., 2007). Briefly, the p3121 plasmid, carrying luciferase was used as a template and amplified with target gene-specific primers. PCR products were then purified, and the residual template plasmid was removed by a DpnI restriction digest. The resulting PCR product was analyzed by agarose gel electrophoresis and used for electroporation into competent cells of *S*. Typhimurium, harboring pKD46. Proper integration of the reporter cassette was confirmed by colony PCR using sanA-check-seq-for and sanA-check-seq-rev primers and Sanger sequencing. To determine SanA expression during infection, the *S*. Typhimurium *sanA*_RBS_*::luc* strain proceeded in infection assay with the use of iBMDM according to the protocol described above. After lysis with Triton X-100, an equal number of samples were used to determine CFU by dilutional plating, while the rest was collected by centrifugation 3 min, 13,000xg. The pellet was then resuspended in the lysis buffer (100 mM potassium phosphate buffer [pH 7.8], 2 mM EDTA, 1 % [wt/vol] Triton X-100, 5 mg/ml bovine serum albumin, 1 mM DTT, 5 mg/ml lysozyme), incubated 30 min on ice and sonicated. Lysates were then analyzed by the addition of luciferase reagent LAR (Promega) in white microtiter plates using a Tecan plate reader and represented as relative light units (RLU) per 2x10^6^ bacteria.

### 7. Hemagglutinin-based reporter gene assays and Western Blotting

*S*. Typhimurium WT and *ΔsanA* strains were transformed with pFPV25.1GFPmut3.1Kan-*sicA*- 2xHA construct by electroporation according to Sambrook and Russell (Sambrook and Russell, 2006). For assays, transformants were grown O/N at 37°C, 180 rpm. The next day, cultures were diluted to an OD_600_=0.05 and grown in SPI-I-inducing conditions, as described above. At indicated time points, the equivalent of bacteria to OD_600_=0.4 was collected by centrifugation for 4 min at 4°C, 16,100xg (Westermann et al., 2019). For Western Blot analysis, pellets were suspended in 100 µl of loading dye, incubated for 5 min at 95°C, and proceeded with SDS PAGE in a 15 % gel. The separated proteins were transferred using semi-dry transfer (Biorad) onto nitrocellulose and blocked for 1 h at RT with 5 % fat-free dry milk in PBST (PBS supplemented with 0.1 % Tween-20). A 1:1000 dilution of HA-Tag Rabbit mAb (Cell Signalling, C29F4) in PBST was used as a primary antibody, whereas the secondary antibody was Anti-rabbit peroxidase diluted 1:5000 in PBST (Sigma, A6154). The blots were developed with Clarity Western ECL Substrate (BioRad). After the first blotting, the membrane was incubated with 30 % H_2_O_2_ for 20 min at 37°C to inactivate peroxidase activity (Sennepin et al., 2009). Then, the membrane was washed two times with PBST and processed with Western Blot as described above, however with GFP Mouse mAb as a primary antibody diluted 1:1000 in PBST (Cell Signalling, 4B10) and Anti-mouse peroxidase (Dako, P0447) as a secondary antibody diluted 1:5000. The Western Blots were developed with the use of Chemidoc XRS+ and analyzed using Image Lab software (BioRad).

### 8. GFP-based reporter gene assays

The pFC-p*sicA* plasmid, which allows for the detection of GFP under the control of the *sicA* promoter and constitutive expression of mCherry, was constructed by cloning as described above. *S*. Typhimurium WT and *ΔsanA* strains were transformed with constructs by electroporation according to the Sambrook and Russell protocol (Sambrook and Russell, 2006). For assays, transformants were grown O/N at 37°C, 180 rpm. The next day, cultures were diluted to an OD_600_=0.05 and grown in SPI-I-inducing conditions as described above. At the indicated time points, OD_600_ was measured, equivalent to 3x10^8^ bacteria was collected by centrifugation (6000xg, 5 min, room temperature), and washed with PBS. Next, bacteria were resuspended in PBS, fixed for 30 min in 4 % PFA in the dark, and washed 3 times with PBS. Prior to analysis, bacteria were resuspended in PBS and filtered. Bacteria carrying pFCcGi empty plasmid (mCherry constitutive expression; GFP no expression) or empty plasmid induced by arabinose (mCherry constitutive expression; GFP induced expression) were used as negative and positive GFP controls, respectively. A cellular fluorescence was measured on the BD Fortessa II cell analyzer with Diva 8 software (Becton Dickinson, Franklin Lakes, NJ, USA) with a total of 10,000 events of the bacterial population (gated on Forward Scatter (FSC)-H versus Side Scatter (SSC)-H dot plots). GFP-positive cells were gated on mCherry-positive bacteria and double-positive populations were further analyzed using FlowJo.

### 9. Generation of an antibody against SanA

The peptide DHRFKHLYGLHRDHHHD, corresponding to amino acid residues 165–184 of SanA, was synthesized and coupled to keyhole limpet hemocyanin (KLH) by Davids Biotechnologie GmbH, Regensburg, Germany. The KLH-coupled peptide was used as immunogen for the generation of rabbit antiserum which was further purified according to protocol used by the company. The optimal concentration of antibody determined for Western Blotting application was 20 μg/ml.

### 10. Bacteria fractionation

Bacterial cultures were separated at the early stationary growth phase into soluble and membrane fractions by a lysozyme-EDTA-osmotic shock and further, the inner and outer membranes were selectively extracted with Triton X-100 as described previously (Russel and Kazmierczak, 1993). Fractions were resuspended in 30 % SDS with Laemmli buffer (50 mM Tris-HCl pH 6.8, 2 % SDS, 10 % glycerol, 1 % β-mercaptoethanol, 12.5 mM EDTA, 0.02 % bromophenol blue) and proceeded with Western Blotting. Briefly, samples were normalized according to Bicinchoninic acid (BCA) assay (Thermo), separated by SDS-PAGE on 12 % gel, and transferred onto PVDF membrane using a semidry transblot system (Bio-Rad). Next, the membranes were blocked for 1 h at room temperature with 5 % fat-free dry milk in PBST. They were then incubated overnight with a 1:100 dilution of OmpA Rabbit antiserum (as a marker of the outer membrane) and LepB Rabbit antiserum (as a marker of the inner membrane), both of which were gifts from Prof. R. Dalbey, Ohio State University. Additionally, a 1:500 dilution of SanA Rabbit polyclonal antibody in PBST was used. The blots were developed with Clarity Western ECL Substrate (BioRad) with the use of Chemidoc XRS+ and analyzed using Image Lab software (BioRad).

## Results

### SanA is located in the inner membrane

The subcellular localisation of the SanA has not been determined so far, and our knowledge about its location within bacterial compartments is based solely on bioinformatic prediction tools. Furthermore, the results of these analysis are not consistent, indicating that SanA can be an outer or an inner membrane protein (Aleksandrowicz et al., 2024). To explore this aspect, we aimed to analyze the subcellular localisation of SanA in *Salmonella*.

SanA was identified solely in the inner membrane fraction, co-located in this compartment with the control protein OmpA (**Fig. 1**). The absence of SanA expression in the *ΔsanA* deletion mutant confirmed the specificity of the newly created SanA antibody, evidenced by the absence of non-specific binding. Furthermore, the membrane marker proteins LepB (inner membrane) and OmpA (outer membrane) were detected predominantly in their respective fractions (**Fig. 1**). This observation suggests that the cytoplasmic membrane integrity was maintained without significant disruption during the process of fractionation, ensuring the reliability of the experiment results.

**Figure 1.**
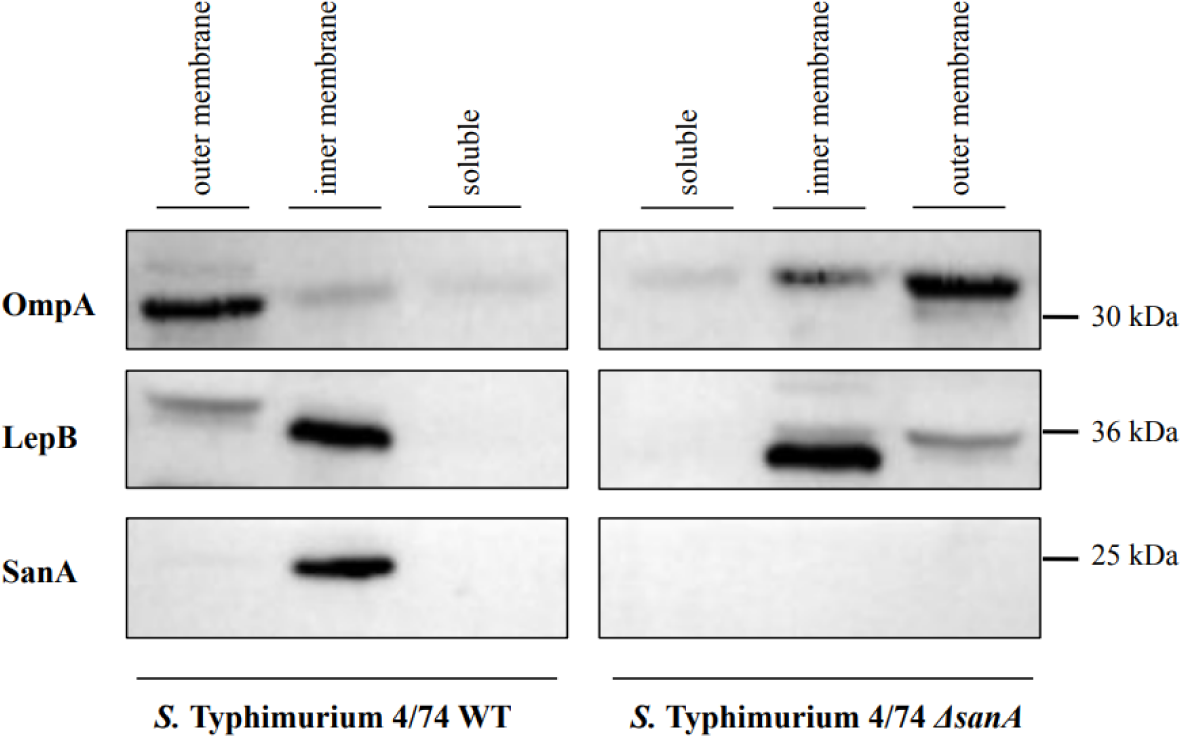
Fractionation of *S*. Typhimurium 4/74 WT and its deletion mutant *ΔsanA* by lysozyme–EDTA method. After fractionation, proteins were subjected to SDS-PAGE. Then, the proteins were blotted onto a PVDF membrane and were probed with the indicated antisera. Proteins from the same number of cells were electrophoresed on each lane. Lane 1, outer membrane fraction; Lane 2, inner membrane fraction; Lane 3, soluble fraction of WT; Lane 4, soluble fraction; Lane 5, inner membrane fraction; Lane 6, outer membrane fraction of *ΔsanA*.

### SanA expression is growth-phase dependent and correlates with intracellular survival within macrophages

To examine the impact of diverse *in vitro* conditions on *sanA* expression, an analysis employing transcriptional fusion was carried out. Addition of transcriptional *luc* reporter had no effect on growth kinetics of the bacteria (**Fig. S1**). We observed strong induction of the reporter in bacteria entering the late exponential and early stationary growth phases (**Fig. 2A**). Only a low level of reporter activity was detected in bacteria during the late stationary growth phase (**Fig. 2A**). Upon culturing bacteria in Mg-MES pH 5.0 medium, resembling conditions in infected macrophages, the activity level was comparable to that in the early exponential growth phase (**Fig. 2A**).

**Figure 2.**
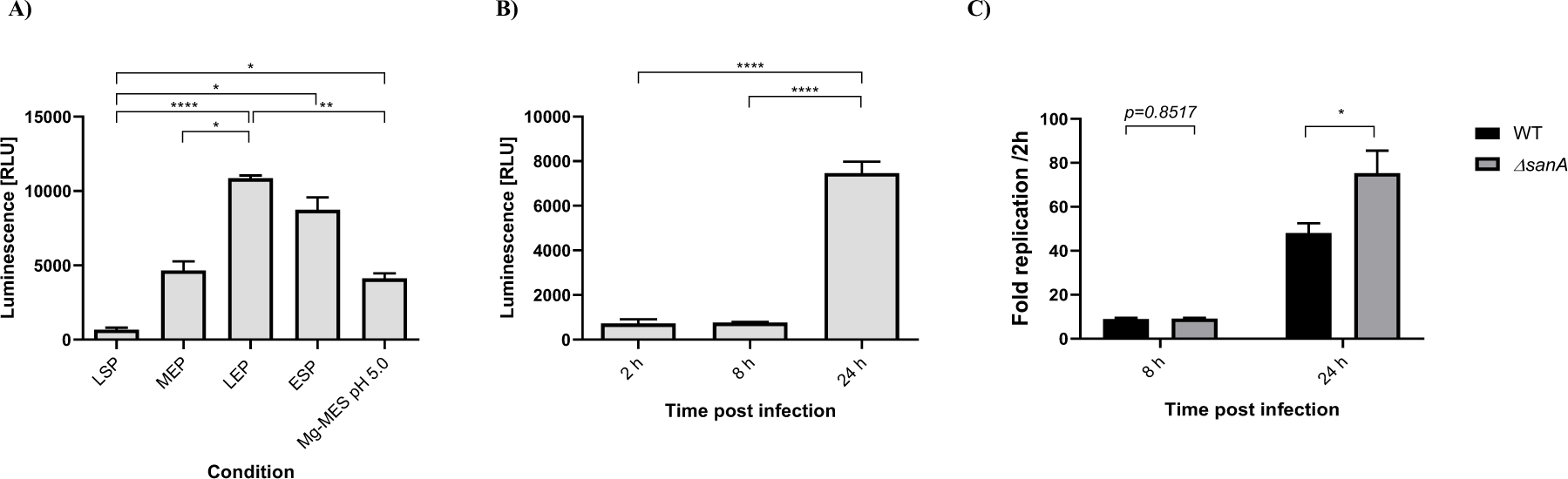
**A)** *In vitro* expression of a *luc* reporter fused to the *sanA* (*sanA*_RBS_*::luc*) in different growth phases; LSP – late stationary phase (16 h culture); MEP – early exponential phase (OD_600_=0.5); LEP – late exponential phase (OD_600_=1.0); ESP – early stationary phase (OD_600_=2.0); Mg-MES pH 5.0 – SPI-II inducing conditions, mimicking environment of infected macrophages. **B)** *In vitro* expression of a *luc* reporter fused to the *sanA* (*sanA*_RBS_*::luc*) during the infection of immortalized BMDM isolated from C57BL/6 mice. C) Intracellular survival within immortalized BMDM isolated from C57BL/6 mice of *S*. Typhimurium 4/74 and its deletion mutant. The data are shown as mean values and SEM of at least three separate experiments. Statistical differences were analyzed by one-way ANOVA with Tukey’s correction (*, p<0.05; **, p < 0.01; ***, p<0.001; ****, p<0.0001).

Similarly, to determine whether *sanA* is significantly expressed at a specific stage following entry into host cells, macrophages were infected with the reporter strain. Subsequently, cells were lysed at different intervals post-infection for the quantification of luciferase activity. The *sanA* expression was low and comparable between 2 h and 8 h post- infection (**Fig. 2B**). The highest expression was detected after 24 h, what was further associated with the differences in intracellular survival of *Salmonella* within immortalized macrophages.

Indeed, we showed no differences in the intracellular bacteria 8 h post-infection and significantly higher replication of the deletion mutant as compared to the WT strain (**Fig. 2C**).

### SanA deletion increases the invasion of intestinal epithelial cells and macrophages

Given prior studies indicating a possible involvement of SanA in the initial stages of infection, we utilized invasion assays to examine this phenomenon (Kolenda et al., 2021). Human epithelial cell line Caco-2 and immortalized bone marrow mice macrophages (iBMDM) were infected with SPI-I-induced *Salmonella* strains at the multiplicity of infection (MOI)=100, and MOI=10, respectively. We noticed that the number of invading WT bacteria was significantly lower than *ΔsanA*, which revealed more than 32 % and about 42 % higher invasiveness towards Caco-2 and iBMDM, respectively (**Fig. 3A** and **3B**). Moreover, the complementation of mutation *in trans* restored the WT phenotype, as *ΔsanA-*pWSK29 invaded both cells type significantly better than *ΔsanA*-pWSK29-*sanA* (**Fig. 3A** and **3B**). Invasiveness towards the Caco-2 cell line was about 16 % higher, whereas in the case of iBMDM 18 % more bacteria invaded the cells.

**Figure 3.**
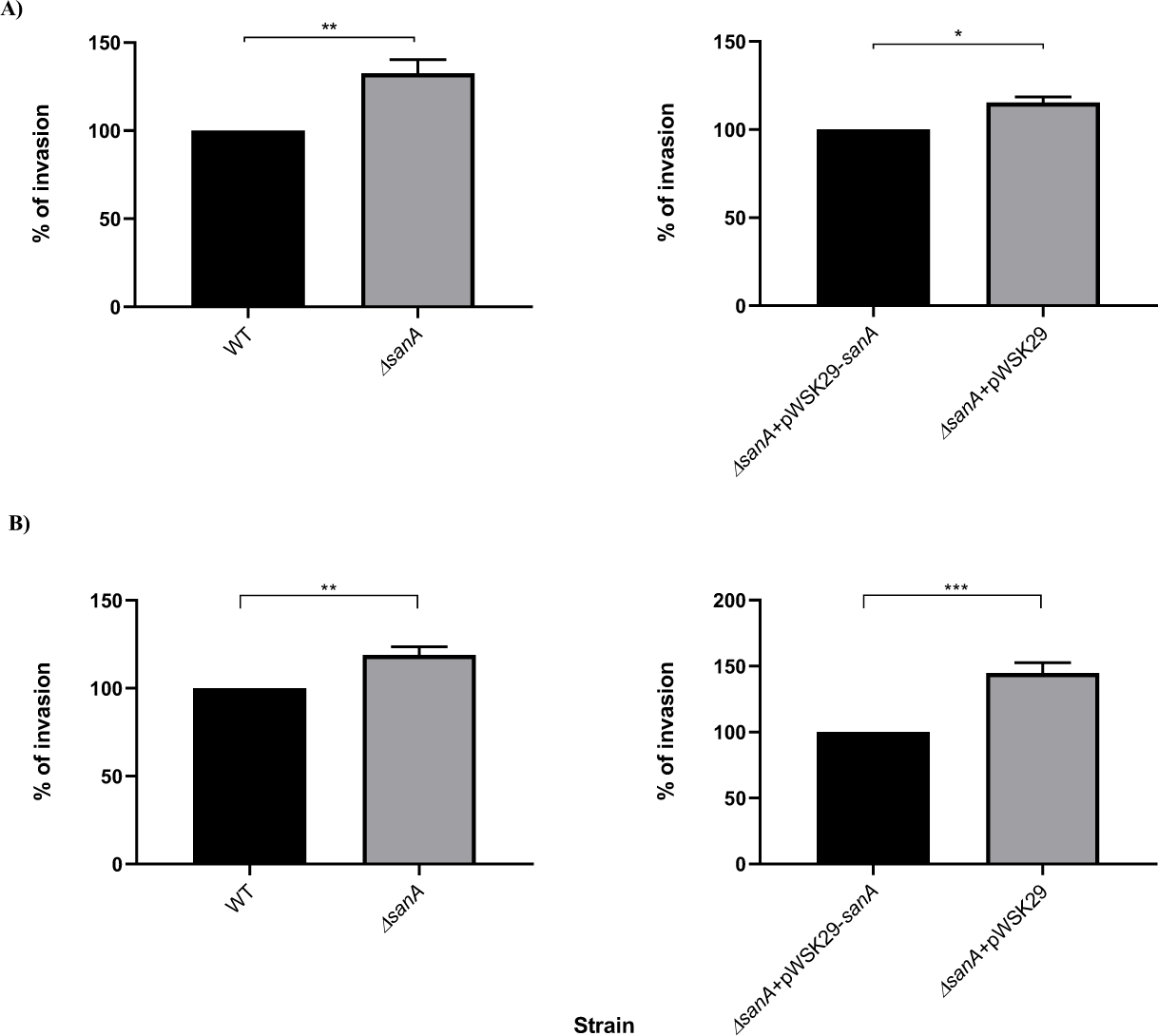
Invasion assay of **A)** Caco-2 cell line and **B)** immortalized bone marrow macrophages isolated from C57BL/6 mice infected by *S*. Typhimurium 4/74, its deletion mutant *ΔsanA* and *ΔsanA* transformed with empty pWSK29 plasmid or vector with *sanA*. The data are shown as mean values and SEM of at least three separate experiments of invasion assay statistical differences were analyzed by Student’s t-test (*, p<0.05; **, p < 0.01).

### SanA deletion correlates with enhanced expression of SicA

As we observed higher invasiveness of *Salmonella* because of a SanA knockout, we decided to analyze the molecular basis responsible for this phenotype. Keeping in mind that SPI-I is the main determinant responsible for the invasion of host cells, our examination included a comparative analysis of type III secretion-associated chaperone SicA promoter activity across populations, and protein expression in the WT and *ΔsanA.* For SicA, the differences were detected in the early stationary growth phase, whereas a very low expression with no differences between analyzed strains was shown in the early exponential growth phase (**Fig. 4A**). These results were examined quantitatively in densitometric analysis, where the average relative density of SicA was assessed against the relative density of GFP, used as a protein load control in the bacterial lysates (**Fig. S2**).

**Figure 4.**
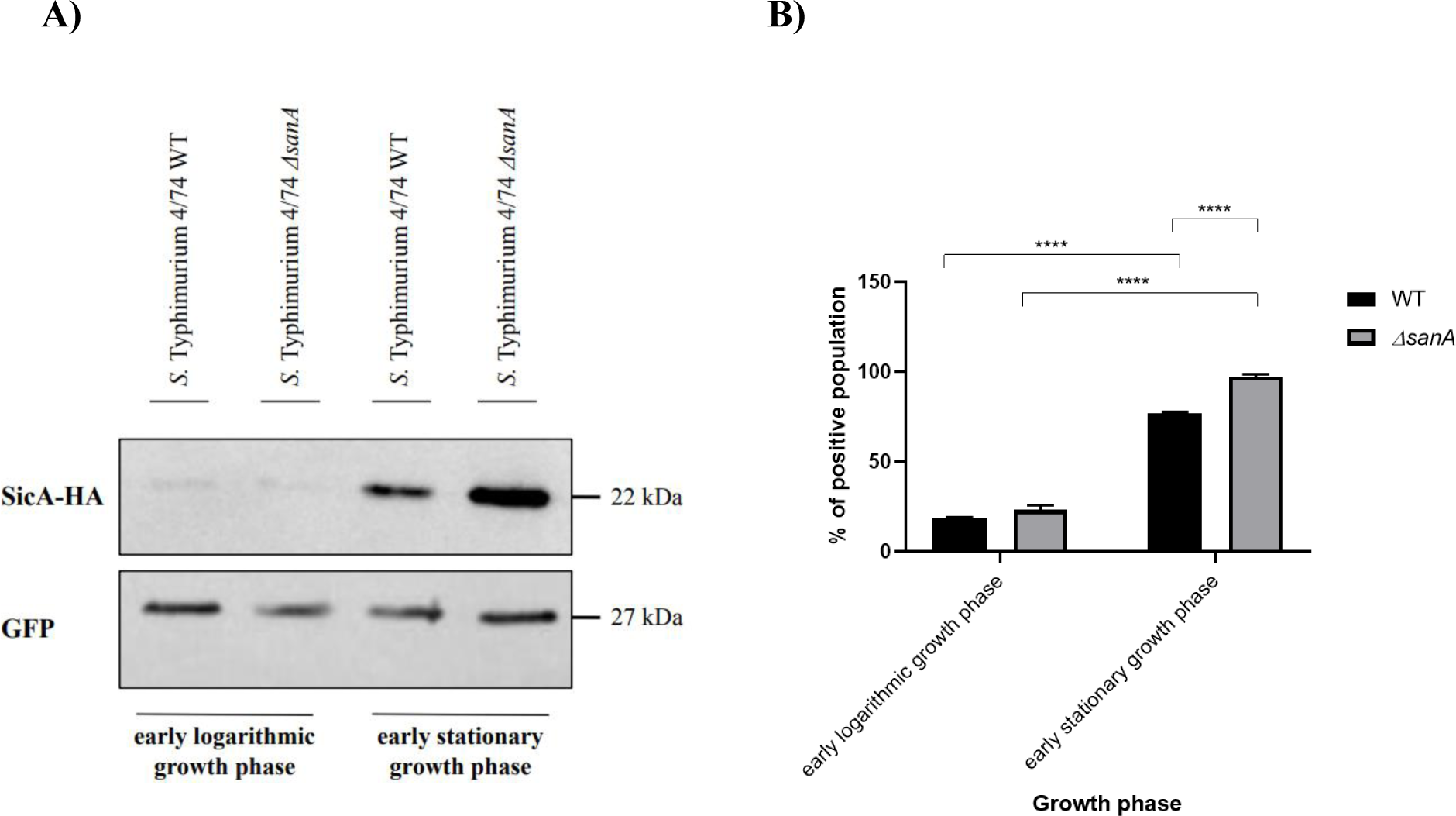
**A)** Determination of SicA expression by Western Blotting; depending on growth conditions in *S*. Typhimurium 4/74, its deletion mutant *ΔsanA*. Early exponential growth phase corresponding to OD_600_=0.5; early stationary growth phase corresponding to OD_600_=2.0; **B)** Fraction of cells expressing *sicA* (on state) during growth in LB medium until early exponential or early stationary growth phase. The fraction of cells in the on state was determined relative to the negative control (100% in the off state), which consisted of the measured fluorescence of cells not expressing the GFP. The data are shown as mean values and SEM of at least three separate experiments. Statistical differences were analyzed by ANOVA (*, p<0.05; **, p < 0.01).

These findings are consistent with our FACS analysis of the reporter system based on the GFP under the control of the promoter of interest and constitutive expression of mCherry. This examination showed a greater proportion of the *ΔsanA* population expressed *sicA*, particularly noticeable during the early stationary growth phase (**Fig. 4B, Fig. S3**). In this phase, approximately 72 % of the WT population and about 95 % of the *ΔsanA* population expressed *sicA* (**Fig. 4B, Fig. S3**).

### SPI-I expression in *ΔsanA* is regulated in a nutrient accessibility dependent manner

The observation of a higher expression of SicA in the *sanA* deletion mutant raised a question of what was responsible for this molecular pattern. As previously demonstrated, high levels of nutrients, resulting from improved accessibility or enhanced transport to bacteria, are associated with heightened virulence in pathogens (Penttinen et al., 2016; Hamed et al., 2019). To address this issue, we employed the abovementioned reporter system to analyse if similar regulation occurs in *S*. Typhimurium. When we grew cells in LB containing 2 % Yeast Extract (YE), we observed the highest (>86 %) population of cells where the *sicA* promoter was active for both, WT and *ΔsanA,* whereas the lowest percentage of on state population was shown with the absence of the yeast extract (<66 %) (**Fig. 5, Fig. S4**). Furthermore, in the presence of 0.5 % YE, there was a shift for *ΔsanA* as compared to WT, as the population of cells where the *sicA* promoter was active was 62 % and 83 %, respectively (**Fig. 5, Fig. S4**).

**Figure 5.**
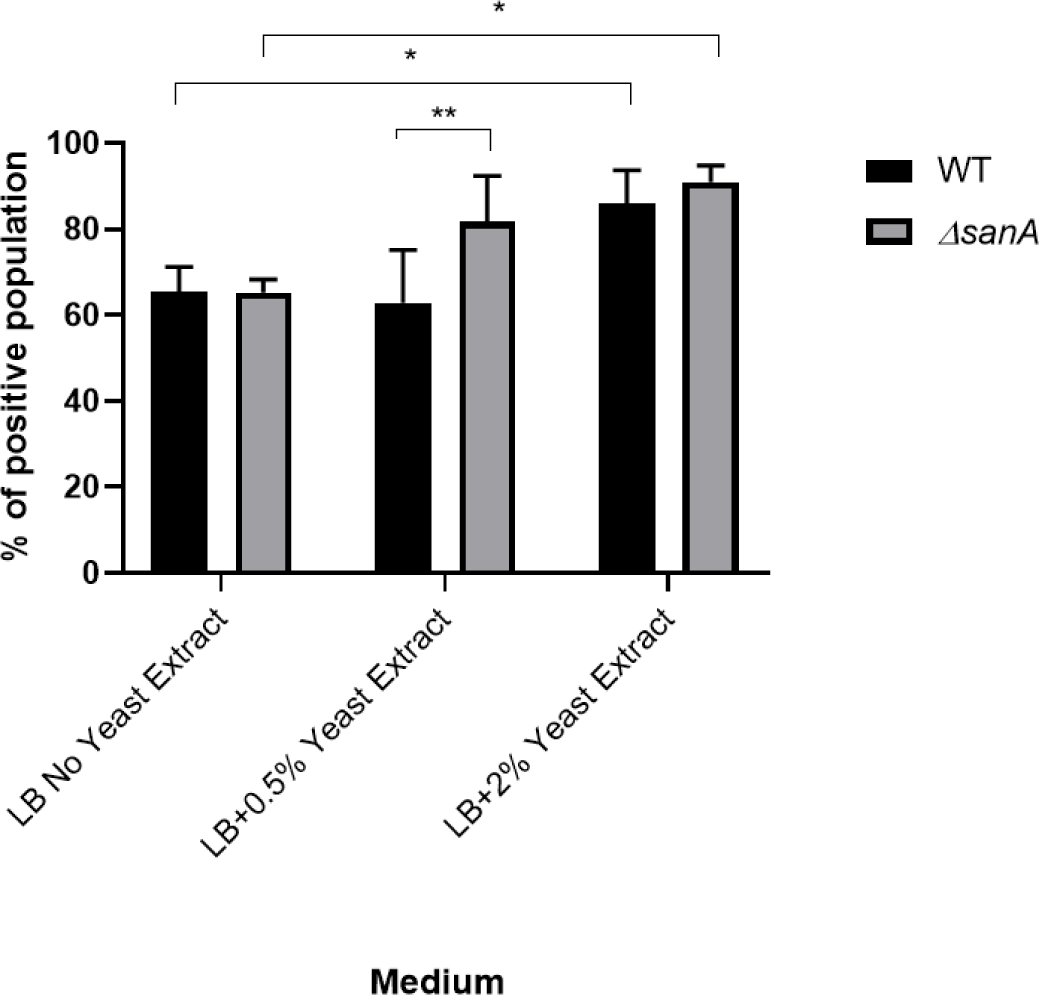
Fraction of cells expressing *sicA* (on state) during growth in LB medium with different concentrations of the yeast extract. The fraction of cells in the on state was determined relative to the negative control (100% in the off state), which consisted of the measured fluorescence of cells not expressing the GFP (without *sicA* promoter). The data are shown as mean values and SEM of at least three separate experiments. Statistical differences were analyzed by ANOVA (*, p<0.05; **, p < 0.01).

## Discussion

Bacterial membranes are composed of numerous proteins that are crucial in the interaction between the pathogen and both the environment and the host (Delcour, 2009; van der Heijden et al., 2016). Among these molecules is SanA, which was first described by Rida et al. in 1996, who identified it as a protein contributing to vancomycin resistance. Specifically, they noted that increased overexpression of SanA reduced the vancomycin sensitivity of *E. coli* mutant with an unidentified envelope permeability defect (Rida et al., 1996). Similarly, *S*. Typhimurium’s SfiX, an ortholog of SanA, was found to mitigate the cell division defect caused by HisHF overproduction (Mouslim et al., 1998).

In our previous studies, we confirmed that SanA is a key player in antibiotic resistance and suggested the association of this phenotype with the physicochemical properties of the bacterial membrane (Aleksandrowicz et al., 2024). Furthermore, we showed that a 10- nucleotide mutation in the *sanA*-encoding sequence results in an enhanced invasion of intestinal epithelial cells of human origin (Kolenda et al., 2021). However, the mechanisms underlying this phenotype remain unexplored. Given the current gaps in understanding SanA’s role in pathogenicity, our objective was to investigate the impact of SanA on the infection phenotype of *S*. Typhimurium 4/74 and explore the correlation between SanA-dependent membrane permeability and host-pathogen interaction.

Although SanA affects the physicochemical properties of the bacterial membrane its subcellular localization remains unverified (Aleksandrowicz et al., 2024). To address this issue, we investigated whether SanA is anchored in the inner membrane and interacts with the components of the outer membrane, thereby inducing specific modifications in this compartment. Alternatively, we considered whether SanA is an outer membrane protein and independently maintains the integrity of the cell envelope. By employing subcellular fractionation and Western blotting techniques, we unequivocally demonstrated that SanA is indeed an inner membrane protein, which aligns with our initial hypothesis. This situation parallels that of TolA, where a mutation in the *tolA* gene results in heightened permeability of the outer membrane. Similar to SanA, TolA is embedded in the inner membrane through its hydrophobic N-terminal segment, which comprises 21 amino acid residues. It is believed that it interacts with components on the inner surface of the outer membrane through its carboxyl-terminal domain to maintain its integrity (Levengood et al., 1991; Levengood-Freyermuth et al., 1993).

To investigate the role of SanA in *Salmonella* infection, we employed two distinct models: human intestinal epithelial cells and immortalized mouse macrophages. These models allowed us to assess invasiveness and intracellular survival, shedding light on SanA’s multifaceted function. Notably, when *Salmonella* was cultured under conditions conducive to the expression of SPI-I, the deletion of *sanA* led to an increase in bacterial invasion of both epithelial cells and macrophages. This finding aligns with our earlier investigation, where we observed enhanced invasiveness due to a 10-nucleotide deletion in *sanA* within a human cell model (Kolenda et al., 2021). Furthermore, this phenomenon correlates with the level of *sanA* expression, as we demonstrated that the protein reached its peak in SPI-I-inducing conditions, particularly during the early stationary growth phase. This observation is consistent with the findings of Kroger et al., who reported a similar expression pattern at the RNA level (Kröger et al., 2013). We propose that the pronounced level of SanA under these conditions is sufficient for the observation of the phenotype, given the substantial difference in expression between the WT and *ΔsanA*. Additionally, in our prior research, we established that the invasiveness level of bacteria remains unchanged when cultivated under conditions that induce the SPI-II (late stationary growth phase), in which the SanA amount is noticeably lower. It suggests a specific association between the phenotype we observed and the correlation between *sanA* expression level and SPI-I (Aleksandrowicz et al., 2024).

Similarly, within the macrophage model, we observed a higher replication rate of *sanA*- deleted bacteria compared to the WT strain after 24 hours of the assay, mirroring the observed pattern of SanA expression during infection. SanA levels increased steadily throughout the assay and were significantly higher after 24 h compared to the 8 h. Moreover, this observation is consistent with outcomes from our previous research, where we demonstrated an analogous pattern in primary bone marrow-derived macrophages. This finding prompted us to hypothesize a correlation between this phenotype, membrane permeability, and the bacterium’s resistance toward host immune response (Aleksandrowicz et al., 2024). The dynamic expression levels of bacterial proteins are known to be influenced by the growth phase and environmental conditions. Significantly, inner membrane proteins like RpoS, AcrAB, FtsH, or SecA are typically expressed at relatively high levels during the early stationary growth phase. These proteins play pivotal roles in adapting to stress, nutrient limitations, and the transition from exponential growth to the stationary phase (Fischer et al., 2002; Hsieh et al., 2011; Mitchell et al., 2017; Kapach et al., 2020). Coupled with earlier research from 1998, which indicated that *sanA* deletion inhibits cell division only at temperatures of 43°C or higher, it becomes increasingly plausible that SanA is vital for bacterial adaptation to harsh environments and is significantly expressed in the stress conditions (Mouslim et al., 1998). The multifaceted role of SanA in *Salmonella* infection emphasizes its significance in the pathogenicity and adaptive response of these bacteria.

Keeping in mind the observed phenotype, our subsequent aim was to elucidate the regulatory dynamics of virulence genes following the host cells invasion. Importantly, our previous genome sequencing data revealed that strain harboring a mutated *sanA* gene displays the gene expression profiles characteristic of highly infectious strains (Kolenda et al., 2021). Therefore, our focus shifts to *sicA*, a crucial component of the *sic-sip* secreted-effector operon within SPI-I known for its essential role in initiating infection. SicA’s expression pattern is expected to closely mirror that of *hilA*, a central regulator of SPI-1 genes (Hensel et al., 1998; Deiwick and Hensel, 1999; Chakravortty et al., 2002). To delve into this, we utilized a GFP transcriptional reporter and investigated its expression patterns in the presence or absence of *sanA*. Remarkably, our findings revealed that SPI-1 gene expression was notably up-regulated in *ΔsanA* during the early stationary growth phase, the SPI-I inducing conditions, under which our infection assays were conducted. This observation strongly suggests a direct link between these regulatory dynamics and the presence of SanA. Intriguingly, a similar outcome was reported in 2020 by Kirthinka et al., who highlighted the role of the Lon protease in controlling SPI-1 genes across a range of stress conditions (Kirthika et al., 2020). As previously hypothesized, a high expression of virulence factors may disrupt the integrity of the bacterial membrane (Bustamante et al., 2008). This observation raises an intriguing possibility: the overexpression of T3SS, resulting from *sanA* deletion, may lead to higher membrane permeability. This scenario is consistent with antibiotic resistance mechanisms and the overall infection phenotype, where membrane permeability can significantly influence bacterial survival and virulence. Virulence gene expression can impose a significant fitness cost on pathogenic bacteria (Sobota et al., 2022). Importantly, Sobota et al. discovered that the expression of *hilD*, a key regulator of *Salmonella* virulence genes, can increase membrane permeability and render bacteria more susceptible to stresses that disrupt the bacterial envelope (Sobota et al., 2022). Therefore, it is compelling to conclude that the heightened membrane permeability observed in the mutant strain, coupled with the overexpression of SPI-I in *ΔsanA* is a direct consequence of these interrelated events. This tight regulation of bimodal virulence gene expression serves to hinder the fixation of attenuated mutants during infection, ensuring the transmission of the virulent genotype, and being an example of bacterial adaptation and survival strategies (Sobota et al., 2022).

As the cell envelope plays a crucial role in the interaction between bacteria and their host, any changes within this compartment can significantly impact the infection phenotype. One notable alteration in the cell envelope is enhanced permeability, which allows for more efficient transport of essential nutrients and ions, supporting bacteria survival and proliferation within host cells (Shimizu, 2013; Hamed et al., 2019). Recent research unveiled intriguing connections between nutrient availability and gene expression in bacteria. For example, it has been observed that nutrients such as yeast extract can induce the expression of the SPI-I gene in *Salmonella* (Hamed et al., 2019, 2021). The yeast extract has been found to induce flagellar gene expression via RflP (also known as YdiV) (Wada et al., 2011). Interestingly, this induction does not appear to be driven by a specific metabolite but rather results from the improved growth facilitated by these nutrients. Although yeast extract’s exact role in invasion remains unknown, it serves as a chemical signal that modulates both SPI-I and flagellar gene expression. This suggests that yeast extract likely acts as a surrogate for other metabolites present in the distal small intestine (Hamed et al., 2021). With these insights in mind, our research aimed to investigate the correlation between enhanced membrane permeability, increased nutrient transport across membranes, and the expression of *sicA* in *Salmonella*. Our findings demonstrated that *sicA* expression was dependent on the level of yeast extract present in the LB medium for both analyzed *Salmonella* strains. Moreover, we observed a significantly higher content of GFP-positive *ΔsanA* cells when 0.5 % yeast extract was present, a condition like the environment where bacteria grew for infection assays in our experimental setup. Based on these results, we can conclude that increased invasiveness is at least partly a consequence of enhanced SPI-I expression, which, in turn, is regulated in a nutrient-accessibility dependent manner.

Taken together, we conclude that the inner membrane protein SanA is of significance for *Salmonella* stress environment response regulation, which is an important aspect of both *Salmonella* entry and survival in macrophages and the gastrointestinal tract. We suggest that SanA is a mediator of virulence genes hosted in the SPI-1 genomic region. This study adds insight into the fate of *Salmonella* in the absence of SanA and highlights the importance of inner membrane proteins in the context of bacteria pathogenicity, especially in the early stages of infection.

## Bibliography

1. Aleksandrowicz, A., Kolenda, R., Baraniewicz, K., Thurston, T. L. M., Suchański, J., and Grzymajlo, K. (2024). Membrane properties modulation by SanA: implications for xenobiotic resistance in Salmonella Typhimurium. Front Microbiol 14. doi: 10.3389/fmicb.2023.1340143.

2. Boughner, L. A., and Doerrler, W. T. (2012). Multiple deletions reveal the essentiality of the DedA membrane protein family in Escherichia coli. Microbiology (N Y) 158, 1162–1171. doi: 10.1099/mic.0.056325-0.

3. Bustamante, V. H., Martínez, L. C., Santana, F. J., Knodler, L. A., Steele-Mortimer, O., and Puente, J. L. (2008). HilD-mediated transcriptional cross-talk between SPI-1 and SPI-2. Proc Natl Acad Sci U S A, 105(38): 14591–14596. doi: 10.1073/pnas.0801205105

4. Chakravortty, D., Hansen-Wester, I., and Hensel, M. (2002). Salmonella pathogenicity island 2 mediates protection of intracellular Salmonella from reactive nitrogen intermediates. Journal of Experimental Medicine. 195(9):1155–66. doi: 10.1084/jem.20011547.

5. Coburn, B., Sekirov, I., and Finlay, B. B. (2007). Type III secretion systems and disease. Clin Microbiol Rev 20, 535–549. doi: 10.1128/CMR.00013-07.

6. Dam, S., Pagès, J. M., and Masi, M. (2018). Stress responses, outer membrane permeability control and antimicrobial resistance in enterobacteriaceae. Microbiology (United Kingdom) 164, 260–267. doi: 10.1099/mic.0.000613.

7. Deiwick, J., and Hensel, M. (1999). Regulation of virulence genes by environmental signals in Salmonella typhimurium. in Electrophoresis (Wiley-VCH Verlag), 813–817. doi: 10.1002/(SICI)1522-2683(19990101)20:4/5<813::AID-ELPS813>3.0.CO;2-Q.

8. Delcour, A. H. (2009). Outer membrane permeability and antibiotic resistance. Biochim Biophys Acta Proteins Proteom 1794, 808–816. doi: 10.1016/j.bbapap.2008.11.005.

9. Ferrari, R. G., Rosario, D. K. A., Cunha-Neto, A., Mano, S. B., Figueiredo, E. E. S., and Conte- Juniora, C. A. (2019). Worldwide epidemiology of Salmonella serovars in animal-based foods: A meta-analysis. Appl Environ Microbiol 85. doi: 10.1128/AEM.00591-19.

10. Figueira, R., Watson, K. G., Holden, D. W., and Helaine, S. (2013). Identification of Salmonella pathogenicity island-2 type III secretion system effectors involved in intramacrophage replication of S. enterica serovar typhimurium: Implications for rational vaccine design. mBio 4. doi: 10.1128/mBio.00065-13.

11. Fischer, B., Rummel, G., Aldridge, P., and Jenal, U. (2002). The FtsH protease is involved in development, stress response and heat shock control in Caulobacter crescentus. Mol Microbiol 44, 461–478. doi: 10.1046/j.1365-2958.2002.02887.x.

12. Gerlach, R. G., Hölzer, S. U., Jäckel, D., and Hensel, M. (2007). Rapid engineering of bacterial reporter gene fusions by using red recombination. Appl Environ Microbiol. 73(13):4234–42. doi: 10.1128/AEM.00509-07.

13. Hamed, S., Shawky, R. M., Emara, M., Slauch, J. M., and Rao, C. V. (2021). HilE is required for synergistic activation of SPI-1 gene expression in Salmonella enterica serovar Typhimurium. BMC Microbiol 21. doi: 10.1186/s12866-021-02110-8.

14. Hamed, S., Wang, X., Shawky, R. M., Emara, M., Aldridge, P. D., and Rao, C. V. (2019). Synergistic action of SPI-1 gene expression in Salmonella enterica serovar typhimurium through transcriptional crosstalk with the flagellar system. BMC Microbiol 19, 211. doi: 10.1186/s12866-019-1583-7.

15. Hensel, M., Shea, J. E., Waterman, S. R., Mundy, R., Nikolaus, T., Banks, G., et al. (1998). Genes encoding putative effector proteins of the type III secretion system of Salmonella pathogenicity island 2 are required for bacterial virulence and proliferation in macrophages. Mol Microbiol 30, 163–174. doi: 10.1046/j.1365-2958.1998.01047.x.

16. Hsieh, Y. H., Zhang, H., Lin, B. R., Cui, N., Na, B., Yang, H., et al. (2011). SecA alone can promote protein translocation and ion channel activity: SecYEG increases efficiency and signal peptide specificity. Journal of Biological Chemistry 286, 44702–44709. doi: 10.1074/jbc.M111.300111.

17. Kapach, G., Nuri, R., Schmidt, C., Danin, A., Ferrera, S., Savidor, A., et al. (2020). Loss of the Periplasmic Chaperone Skp and Mutations in the Efflux Pump AcrAB-TolC Play a Role in Acquired Resistance to Antimicrobial Peptides in Salmonella typhimurium. Front Microbiol 11. doi: 10.3389/fmicb.2020.00189.

18. Kirthika, P., Senevirathne, A., Jawalagatti, V., Park, S. W., and Lee, J. H. (2020). Deletion of the lon gene augments expression of Salmonella Pathogenicity Island (SPI)-1 and metal ion uptake genes leading to the accumulation of bactericidal hydroxyl radicals and host pro-inflammatory cytokine-mediated rapid intracellular clearance. Gut Microbes 11, 1695–1712. doi: 10.1080/19490976.2020.1777923.

19. Kolenda, R., Burdukiewicz, M., Wimonc, M., Aleksandrowicz, A., Ali, A., Szabo, I., et al. (2021). Identification of Natural Mutations Responsible for Altered Infection Phenotypes of Salmonella enterica Clinical Isolates by Using Cell Line Infection Screens. Appl Environ Microbiol, 87(2):e02177–20. doi: 10.1128/AEM.

20. Kröger, C., Colgan, A., Srikumar, S., Händler, K., Sivasankaran, S. K., Hammarlöf, D. L., et al. (2013). An infection-relevant transcriptomic compendium for salmonella enterica serovar typhimurium. Cell Host Microbe 14, 683–695. doi: 10.1016/j.chom.2013.11.010.

21. Levengood SK, Beyer WF, Webster RE. 1991. TolA: A membrane protein involved in colicin uptake contains an extended helical region. National Academy of Sciences. USA. 88(14): 5939–5943. doi: 10.1073/pnas.88.14.5939

22. Levengood-Freyermuth SK, Click EM, Webster RE. 1993. Role of the Carboxyl-Terminal Domain of TolA in Protein Import and Integrity of the Outer Membrane. Journal of Bacteriology. 175(1):222–8. doi: 10.1128/jb.175.1.222-228.1993

23. Martínez, L. C., Banda, M. M., Fernández-Mora, M., Santana, F. J., and Bustamante, V. H. (2014). HilD induces expression of Salmonella pathogenicity Island 2 genes by displacing the global negative regulator H-NS from ssrAB. J Bacteriol 196, 3746–3755. doi: 10.1128/JB.01799-14.

24. Matz, C., and Jürgens, K. (2001). Effects of hydrophobic and electrostatic cell surface properties of bacteria on feeding rates of heterotrophic nanoflagellates. Appl Environ Microbiol, 67, 814– 820. doi: 10.1128/AEM.67.2.814-820.2001.

25. Mitchell, A. M., Wang, W., and Silhavy, T. J. (2017). Novel RpoS-dependent mechanisms strengthen the envelope permeability barrier during stationary phase. J Bacteriol 199. doi: 10.1128/JB.00708-16.

26. Monack DM, Raupach B, Hromockyj AE, Falkow S. 1996. Salmonella typhimurium Invasion Induces Apoptosis in Infected Macrophages Source. Proc Natl Acad Sci U S A. 93(18):9833– 9838. doi: 10.1073/pnas.93.18.9833

27. Mouslim, C., Cano, D. A., and Casadesús, J. (1998). The sfiX, rfe and metN genes of Salmonella typhimurium and their involvement in the His(c) pleiotropic response. Molecular and General Genetics, 259(1):46–53. doi: 10.1007/s004380050787.

28. Penttinen, R., Kinnula, H., Lipponen, A., Bamford, J. K. H., and Sundberg, L. R. (2016). High Nutrient Concentration Can Induce Virulence Factor Expression and Cause Higher Virulence in an Environmentally Transmitted Pathogen. Microb Ecol 72, 955–964. doi: 10.1007/s00248-016-0781-1.

29. Peterson J. W (1996). Bacterial Pathogenesis in Medical Microbiology. 4th edition.

30. Pizarro-Cerdá, J., and Cossart, P. (2006). Bacterial adhesion and entry into host cells. Cell 124, 715– 727. doi: 10.1016/j.cell.2006.02.012.

31. Rida, S., Caillet, J., and Alix, J. H. (1996). Amplification of a novel gene, sanA, abolishes a vancomycin-sensitive defect in Escherichia coli. J Bacteriol, 178(1):94–102. doi: 10.1128/jb.178.1.94-102.1996.

32. Russel, M., and Kazmierczak, B. (1993). Analysis of the Structure and Subcellular Location of Filamentous Phage pIV. J Bacteriol, 175(13): 3998–4007. doi: 10.1128/jb.175.13.3998-4007.1993.

33. Sambrook, J., and Russell, D. W. (2006). Transformation of E. coli by Electroporation . *Cold Spring Harb Protoc*. CSH Protoc, 2006(1):pdb.prot3933. doi: 10.1101/pdb.prot3933.

34. Sennepin, A. D., Charpentier, S., Normand, T., Sarré, C., Legrand, A., and Mollet, L. M. (2009). Multiple reprobing of Western blots after inactivation of peroxidase activity by its substrate, hydrogen peroxide. Anal Biochem 393, 129–131. doi: 10.1016/j.ab.2009.06.004.

35. Shimizu, K. (2013). Regulation Systems of Bacteria such as Escherichia coli in Response to Nutrient Limitation and Environmental Stresses. Metabolites 4, 1–35. doi: 10.3390/metabo4010001.

36. Silhavy, T. J., Kahne, D., and Walker, S. (2010). The bacterial cell envelope. Cold Spring Harb Perspect Biol, 2(5):a000414. doi: 10.1101/cshperspect.a000414.

37. Simpson, B. W., and Trent, M. S. (2019). Pushing the envelope: LPS modifications and their consequences. Nat Rev Microbiol 17, 403–416. doi: 10.1038/s41579-019-0201-x.

38. Sobota, M., Ramirez, P. N. R., Cambré, A., Rocker, A., Mortier, J., Gervais, T., et al. (2022). The expression of virulence genes increases membrane permeability and sensitivity to envelope stress in Salmonella Typhimurium. PLoS Biol 20. doi: 10.1371/journal.pbio.3001608.

39. Sun, J., Rutherford, S. T., Silhavy, T. J., and Huang, K. C. (2022). Physical properties of the bacterial outer membrane. Nat Rev Microbiol 20, 236–248. doi: 10.1038/s41579-021-00638-0.

40. van der Heijden, J., Reynolds, L. A., Deng, W., Mills, A., Scholz, R., Imami, K., et al. (2016). Salmonella rapidly regulates membrane permeability to survive oxidative stress. mBio 7. doi: 10.1128/mBio.01238-16.

41. Wada, T., Morizane, T., Abo, T., Tominaga, A., Inoue-Tanaka, K., and Kutsukake, K. (2011). EAL domain protein YdiV acts as an anti-FlhD4C2 factor responsible for nutritional control of the flagellar regulon in Salmonella enterica serovar typhimurium. J Bacteriol 193, 1600–1611. doi: 10.1128/JB.01494-10.

42. Westermann, A. J., Venturini, E., Sellin, M. E., Förstner, K. U., Hardt, W. D., and Vogel, J. (2019). The major RNA-binding protein ProQ impacts virulence gene expression in salmonella enterica serovar typhimurium. mBio 10. doi: 10.1128/mBio.02504-18.

