## Supplementary material for "SanA is an inner membrane protein mediating the early stages of Salmonella infection"

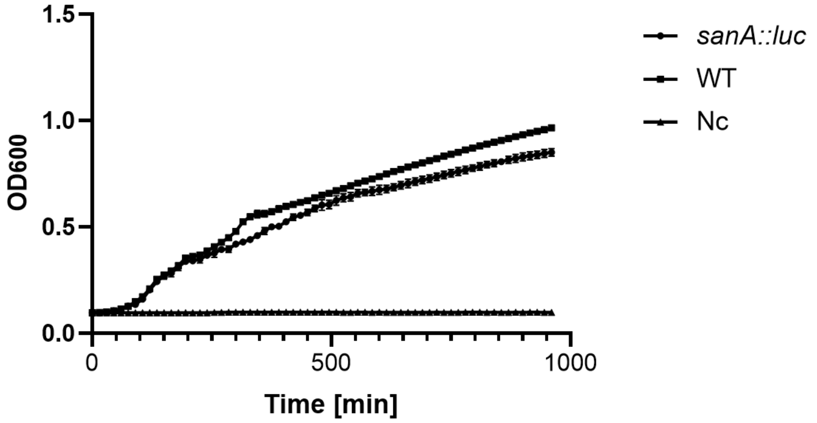


**Figure S1** Growth curves of *S*. Typhimurium 4/74 WT and *sanA*$_{RBS}$*::luc* in LB medium. Nc indicates medium without bacteria. The data are shown as mean values and SEM of at least three separate experiments.

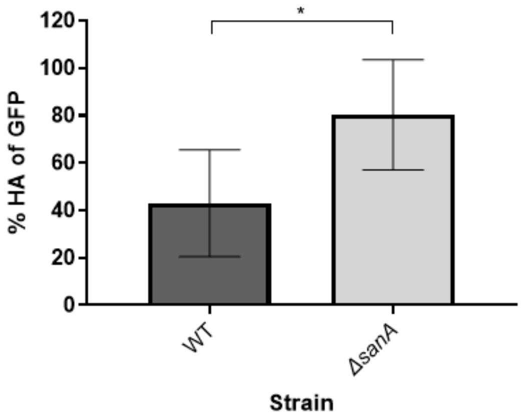


**Figure S2** Densitometric analysis of protein bands imaged with the ChemiDoc MP. The average relative density of SicA was compared to the relative difference in GFP quantity of protein load for the *S*. Typhimurium lysates. The data are shown as mean values and SEM of at least three separate experiments. Statistical differences were analyzed by Student's t-test (*, $p<0.05$; **, $p < 0.01$).

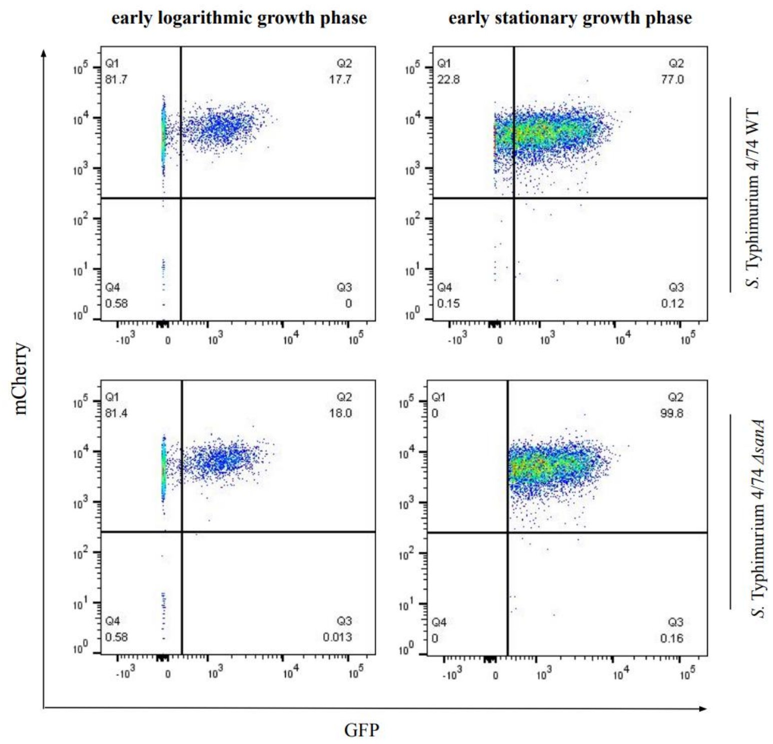


**Figure S3** Fraction of cells expressing *sicA*. The fraction of cells in the on state was determined relative to the negative control (100% in the off state), which consisted of the measured fluorescence of cells not expressing the GFP. Early logarithmic growth phase corresponding to $OD_{600}$=0.5; early stationary growth phase corresponding to $OD_{600}$=2.0.

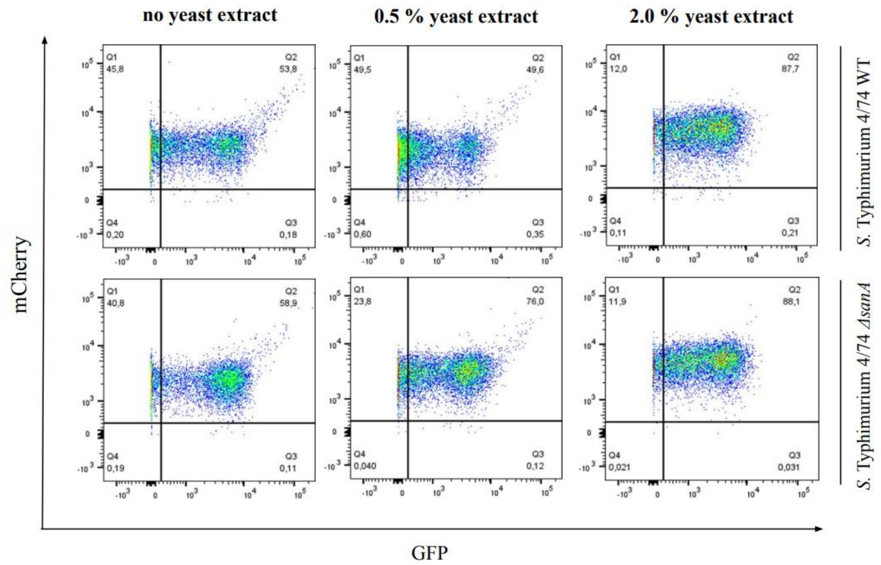


**Figure S4** Fraction of cells expressing *sicA*. The fraction of cells in the on state was determined relative to the negative control (100% in the off state), which consisted of the measured fluorescence of cells not expressing the GFP.
